# Tail Flaring During Agonism in Hummingbirds

**DOI:** 10.64898/2026.01.30.702386

**Authors:** Rosalee Elting, Md Zafar Anwar, Donald R. Powers, Bo Cheng, Haoxiang Luo, Bret W. Tobalske

## Abstract

Bird tails in flight generate lift, reduce pressure drag, and stabilize pitch. Hummingbirds can sustain hovering in still air and generate lift in both the up- and downstroke due to their proportionally enormous primary flight muscles and derived wing skeleton. Given their mass-specific wing power, it is possible that tails are less essential to the aerodynamics of hummingbird flight than they are in other birds, freeing them for non-locomotor functions. Hummingbird tails are well known for their morphological elaboration as sexually selected ornaments, including sound generation. From informal observations of agonism, we hypothesized tail flaring may also be a visual signal used by males during fighting. To begin testing this, we measured the frequency and magnitude of tail flaring during fighting. We measured kinematics of tail flaring during male-male fighting of Calliope hummingbirds (*Selasphorus calliope*, n = 10) indoors. Consistent with our hypothesis, captive males exhibited greater angles of tail flare when engaged in a fight (34.6 ± 45.5°, mean ± sd) than when performing solitary landing (−14.2 ± 9.12 °) and takeoff (−14.1 ± 7.0 °) maneuvers. When birds won a fight, the angles of their tail spread were greater than when they lost (p = 0.01). Moreover, recordings of seven species of hummingbird in the field revealed 95% of inter- and intra- specific contests included tail flaring. We evaluate these results in the context of signaling during animal contests and propose future tests of whether tail flaring is an honest signal of individual quality and Resource Holding Potential.

**Summary Statement:** Male-male fighting is common in hummingbirds with competition over food and mates. During these competitions, tail flaring and waggle maneuvers are used as a signal of aggressive intent.

## Introduction

Self-propelled flight requires more power output than any other form of animal locomotion (Alexander, 2005). Animals that fly must generate sufficient power to remain airborne (weight support), as well as to overcome drag on both the body and the lift-generating surfaces used in flight. While birds’ wings have received the most attention, research spanning the last several decades has suggested several ways the avian tail also serves as a lift-generating surface. To address the demands of weight support, incoming air travels over the tail’s surface and generates lift (Tobalske, 2022). The presence of a tail also provides a more fusiform shape to the bird, serving as a splitter plate and thereby reducing pressure drag on the body (Maybury et al., 2001). How well the tail accomplishes these tasks depends on its shape. Mathematical models of the aerodynamics of delta wings suggest that tails whose widest points are at their trailing edges are the best at maximizing lift relative to drag (Thomas, 1997). Selective pressures on tail shape can address performance flight metrics beyond lift. It is hypothesized that deeply forked tails, instead of maximizing lift to drag ratios, improve maneuvering flight performance by maximizing aerodynamic turning moments (Thomas and Balmford, 1995). A single tail shape can modify morphology to address demands within a wingbeat. In the Warbling White-eye (*Zosterops japonicus*), a passerine species, the tail is flared in mid-downstroke and interacts with wing downwash to mitigate downward pitch of the body (Su et al., 2012). In short, there is evidence that the tail performs diverse aerodynamic roles in volant birds.

Hummingbirds are the only avian species that can sustain hovering in still air, they are extremely agile fliers (Badger et al., 2023; Cheng et al., 2016), and their hovering and maneuvering abilities appear to be due to their small size and specializations of their wings. These specializations include high wingbeat frequencies (20-50 Hz) (Corben, 1983), generation of weight support in both up and down strokes (Warrick et al., 2005), and high mass-specific power output from their proportionally large primary flight muscles (Agrawal et al., 2022) the pectoralis and supracoracoideus, which together account for ∼27% of total body mass (Altshuler et al., 2010; Hartman, 1954). Compared to other avian species, hummingbirds are considered to be “hyper-aerial,” and this lifestyle is hypothesized to have resulted in an evolutionary tradeoff between developmental investment in the forelimbs versus hindlimbs (Heers and Dial, 2015).

This is manifest in hummingbirds having proportionally tiny hindlimbs and using their wings to provide 41% of the initial thrust for takeoff compared to <10% in other birds studied (Earls, 2000; Heppner and Anderson, 1985; Tobalske et al., 2004).

The tail may play an aerodynamic role in hummingbird flight but is better known for its role in sexual selection. An aerodynamic study regarding the tail focused on performance in escape, a dramatic maneuver that requires high power output and agility (Cheng et al., 2016). For the duration of an escape from a looming stimulus, the tail flares constantly; however, computational fluid dynamic modeling of the tail suggests that generation of torque is brief (8% of total time), and peak magnitude is only a fraction of that provided by the wings (25%) (Haque et al., 2023b). While the tail’s role in escape therefore is not nominal, this function seems minor compared to that of mate attraction. Across the hummingbird clade, tails exhibit diverse and fascinating forms, with morphologies including long streamers and spatulate tips, as well as feathers which generate sounds (sonations) in courtship (Clark, 2010). These specialized morphologies may have arisen through decreased evolutionary pressure on the tail for aerodynamic performance (Clark, 2010). This hypothesis has been supported by studies of forward flight, wherein hummingbirds flying in a wind tunnel with experimentally shorter or longer tails suffered minimal reductions in top flight speed and only marginal increases in metabolic rates (Clark and Dudley, 2009).

There is also evidence that the tail is used in hummingbirds’ extremely agile and aggressive territorial defense. Many species of hummingbirds, especially those of the bee clade (McGuire et al., 2014), fiercely defend territories (Dearborn, 1998; Kodric-Brown and Brown, 1978; Nuñez-Rosas et al., 2017; Powers, 1987; Stiles and Wolf, 1970). Descriptions of agonism in several species during territorial defense note the fanning of tails (Hunter, 2008; Hunter and Picman, 2005; Hurly et al., 2001; Williamson, 2020). Herein we use the term “flaring” which we interpret as equivalent to “fanning” and define flaring as an increase in the angle formed between the outer tail rectrices, R5. Territories are worth defending as they provide a source of food and a stage from which to perform impressive courtship dives, shuttles, sonations, and gorget signaling to potential mates (Clark et al., 2018; Hogan and Stoddard, 2018). Defense of the territory from inter- and intraspecific competition can include displacements and chases (Tamm et al., 1989), escalating to feather-pulling or jousting (Rico-Guevara and Araya-Salas, 2015).

While competition seems dramatic and extreme, injury and lethality appear to be rare (Baltosser and Russell, 2020; Evens and Harper, 2020); in other groups of fighting animals, this rarity of injury is the result of preliminary signaling or assessment stages in contests (Hardy and Briffa, 2013; Hoem et al., 2007). Aspects of combat have been documented in several hummingbird species (Bribiesca et al., 2019; Ewald, 1985; González-Gómez et al., 2014; López-Segoviano et al., 2018; Márquez-Luna et al., 2022). Yet open questions remain about how these contests escalate, how individuals communicate their quality or ability to win, and whether tail flaring functions as a signal.

In informal observations of males fighting around outdoor feeders, we noticed frequent use of tail flaring. We hypothesized there may be a difference in the posture of the tail during solitary behaviors compared to competitive interactions. We predicted that tails would be flared at greater angles when in competition, with an alternative prediction that if flaring of the tail was a byproduct of maneuvering in hummingbirds, we would see similar angles in solitary maneuvers (take-off and landing) and competition. We tested these predictions in both laboratory and field settings. To measure details of tail kinematics, we studied Calliope hummingbirds (*Selasphorus calliope)* engaged in agonistic encounters in an indoor flight arena. To place tail flaring in a natural context, we measured the frequency of tail flaring among seven species of wild hummingbirds engaged in fighting outdoors. These two datasets would support our predictions if tail flaring was greatest when in competition and tail flaring was regularly seen in naturally occurring interactions. We would then have preliminary evidence that the tail may serve a communication role during agonism, much like it does in courtship.

## Materials and Methods

### Kinematic Analysis

We obtained videos for 3D kinematic analysis using synchronized cameras. In laboratory studies, we used six cameras: two Nova S6, one Mini AX100, and three Mini UX100 (Photron Inc., San Diego, CA, USA), sampling at 2000 frames s^-1^ with a shutter speed of 1/5000 s. In the field we used three video cameras (two Photron Nova S6 and one MiniAX-100), sampling at 1000 frames s^-1^ with a variable shutter speed dependent on ambient light conditions. We used DLTdv8 (Hedrick, 2008) and EasyWand5 (Theriault et al., 2014) in MATLAB (Mathworks, Inc., 2025) to digitize and calibrate the videos via direct linear transformation to resolve global 3D coordinates.

### Bird Capture and Housing

Male Calliope hummingbirds (*Selaphorus calliope*, 2.6±0.1g, mean ± standard deviation, n = 10) were captured upon arrival at a breeding ground (Year 1: May 14-20, 2024 and Year 2: May 14-15, 2025) in Missoula County, Montana, USA. We trapped birds on private land using a Russell-style drop trap (Russell and Russell, 2001; Tell et al., 2021), then transported them to the Field Research Station at Fort Missoula, Montana, USA. Birds were housed in a 0.6 m^3^ cages constructed of 1mm wire mesh, equipped with feeders providing ad libitum a 50:50 mix of NektarPlus (NEKTON® Günter Enderle, Pforzheim, BadenWürttemberg, Germany) and 20% sucrose (mass: volume). Birds experienced natural lighting and photoperiods during captivity.

Capture and housing were permitted by Montana Fish, Wildlife and Parks (2024-019-W and 2025-030-W) and the US Fish and Wildlife Service (SCCL-771277). Animal procedures were approved by the Institutional Animal Care and Use Committees of the University of Montana (011-23, 055-23). We measured body mass using a digital balance (±0.001g), and projected surface areas of the frontal view of the body and dorsal view of the wings and tail (mm^2^) using digital photographs (Apple IPhone 8se, 7 megapixel, Apple, Inc, Cupertino, CA, USA). Birds were released unharmed after 4-6 weeks of captivity.

### Competition Arena

We constructed a competition arena (1.45m w × 2.9m l × 1.2m h) from aluminum supports and white diffuser fabric. This arena contained a single food source and two perches of differing heights above the floor of the arena (101cm, 71.9cm) to serve as a resource and elicit aggressive interactions (Gaffney, 2017). To measure activity patterns and body weight, we equipped the perches with load cells (Futek S-Beam Jr Load Cell, 10- and 20-gram limits, Futek Advanced Sensor Technology, Inc., Irvine, California, USA). Birds were acclimated individually in the arena. Competition trials only began after birds regularly used perches and regularly fed. During acclimation time, we obtained video recordings of perch landings and takeoffs when birds were calmly completing these maneuvers, without any startle response due to researcher movement. Competition trials occurred between dyads of randomly selected males. Birds were allowed to freely interact for the first 90 minutes of a trial, after which we periodically startled the birds off their perches using hand waving that was visible to the birds via the access door or clear windows in the walls of the arena. This upset the residency (Alcock and Bailey, 1997; Davies, 1978) of the desirable high perch, and required birds to be aerial. The time intervals in which the startles occurred were not measured in this study, we only sampled the subsequent interactions between birds while they were attempting to secure the high perch.

### 3D Kinematic analysis of body positions

The competition arena was surrounded by six synchronized high-speed cameras and was illuminated using LED lights. We digitized ten anatomical landmarks on the birds (beak tip and base and left and right: shoulders, wingtips, tail bases, and tail tips). One out of every 50 frames (2%) were hand-digitized in DLTdv8, and we used these as a training set for Deep Neural Network Learning (DeepLabCut) (Mathis et al., 2018) to digitize remaining frames. We inspected the results for variation in a fixed morphological measure (bill length). If the bill was not visible (obstructed by other bird), we quantified variation only from frames where the bill was visible. When error exceeded 5%, we used hand digitizing to replace the results from DeepLabCut (coefficients of variance for data reported in Table S1).

To measure tail movement, we defined two position vectors, one on the left and the other on the right, originating at the left and right lateral margins of the rump (where the tail meets the body) and moving to the outermost visible left or right rectrix. We obtained the angle of tail flare using the dot product of these vectors. Negative angles represented a tail which was folded, where the distance between the tips of the two rectrices was less than the distance between the left and right rump points (see Fig. 1A for example of a negative tail angle). For each hummingbird (n=10) we measured flaring throughout one takeoff, landing, contest win, and contest loss. To standardize digitized contests, we chose those in which the incoming bird was visible to their competitor (frontal approach). Fights ranged from 0.3 to 3.6 seconds, with an average duration of 0.8 ± 0.5 seconds. To avoid bias due to interval duration, we downsampled measurements of the tail in each individual per flight event in longer events to 612 samples (306 ms) via systematic decimation, selecting every nth point based on each flight’s total duration. To measure bird-centered body rotations (yaw, pitch and roll, using a right-hand coordinate system, Fig. S1), we defined the center of the caudal spine as the average x, y, z coordinates of the base of the tail (rump). The body plane was then defined by two position vectors originating at the rump and moving to the left and right shoulders. We defined the z axis as the cross-product of these two vectors. The bird-centered x axis was from the origin to the center of the spine between the shoulders, and the y axis was the cross product of the x and z axes. We measured yaw about the z axis, pitch about y axis and roll about the x axis.

**Figure 1.**
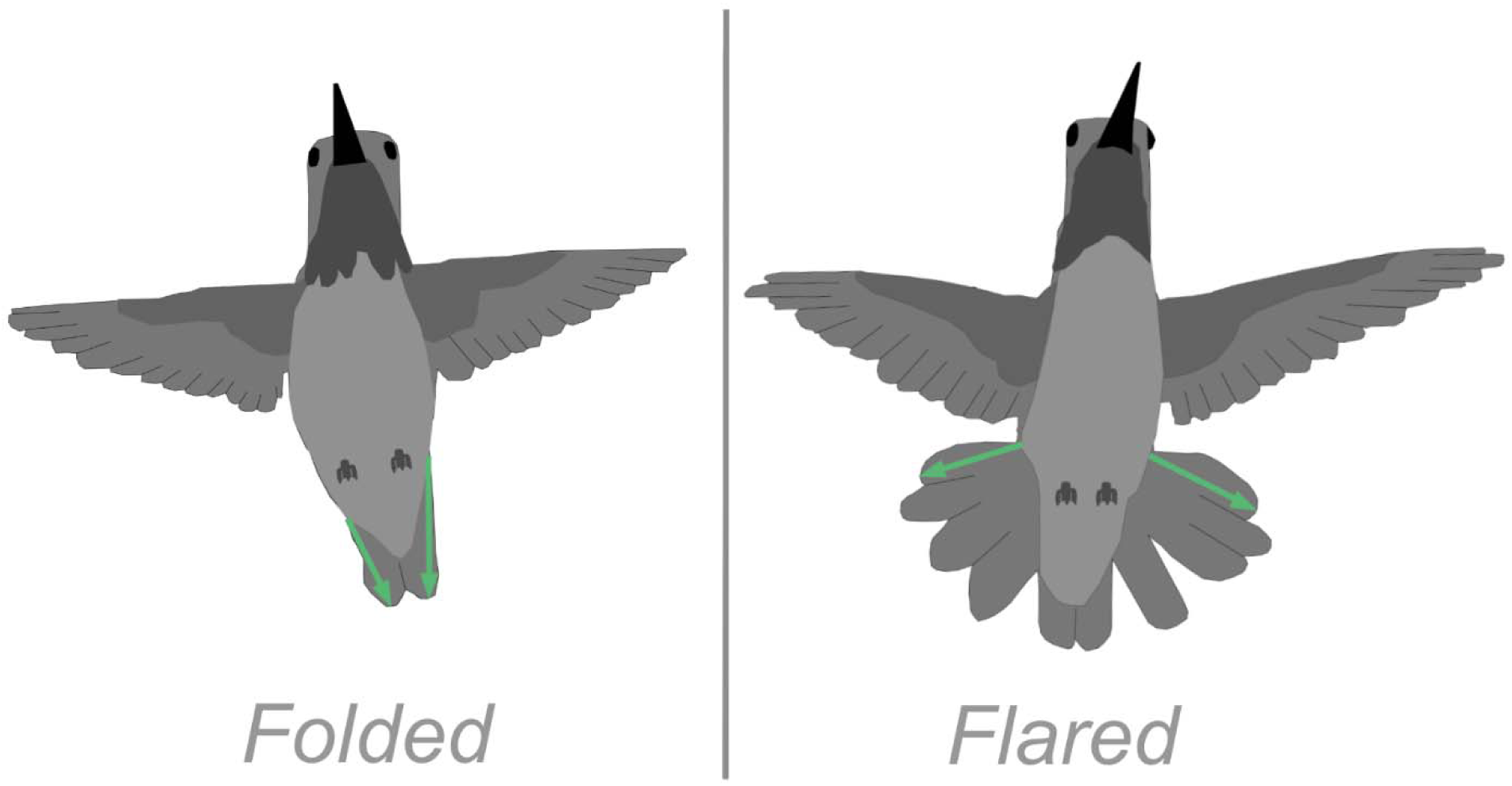
Two tail states in male Calliope hummingbirds (*Selasphorus calliope*): folded and flared. Tail angle was the acute angle between left and right position vectors (green arrows) originating where the outer margin of the tail met the body and going to the tip of each outer rectrix (R5). In these examples drawn from frames of high-speed video, the folded tail angle was −26° and flared angle was 136°.

### Qualitative description of contests

For fights in the competition arena, we watched all contest videos (n=213) to characterize flight trajectories and whether either individual exhibited tail flaring (defined as approximately >30° and sustained for at least 5 ms). Our qualitative visual assessment was informed by our prior 3D measures of tail flaring, described above.

We also recorded when a bird successfully displaced their competitor from the high perch (win) and when a bird retreated (loss). When an incoming bird intimidated their competitor, we recorded if the bird approached from caudal or from the lateral side of the receiving bird, as opposed to the front. From this we analyzed if signaling (tail flaring) effort varied across these approach angles.

### Statistical analysis

The distributions of tail flare angles were heavily skewed with peak values centered around zero. All our statistical tests are based on a single value for each individual (n=10). Because data were not normally distributed, we used a series of non-parametric models to test for differences in tail angles across treatment (takeoff, landing, competition) and to test for relationships between tail flaring and approach angle. We used R for all tests (R core team, 2025): A **Wilcoxon Signed-Rank Test** was used to test for significant differences in the median values of tail flare between two matched samples for n=10 individuals. When we had an *a priori* hypothesis, a single-tailed test was used. From this test a V statistic was generated, which represented how many of the matched samples had a greater value of tail flaring between the first and the second treatment (sum of the rank differences). We tested if tail angles were different between 1) takeoff and competition (one-tailed, *a priori* prediction of greater angles in competition than takeoff), 2) takeoff and landing (two-tailed, no prior prediction) and 3) between contest winners and losers (two-tailed, no prior prediction). We also used this to test for differences in projected frontal area with and without the tail flared in morphometric photos.

We obtained an effect size for all treatment combinations tested in Wilcoxson Signed-Rank Tests using a **Rank-Biserial Correlation** (Kassambara, 2023), which measured the magnitude of differences between groups from Wilcoxon Signed-Rank tests using Z-scored data. This correlation ranges from 0 (no correlation) to 1 (strongest correlation).

We tested whether approach angles affected the use of tail flaring by measuring a proportion of each approach angle (front, side, caudal) where the tail was flared. To do this we used a **Friedman Test**, this is a non-parametric alternative to a repeated-measures ANOVA, run on the ranks of treatments instead of their mean values (Sheldon et al., 1996). To avoid pseudo replication and bias where we had more measurements from some individuals than others, we calculated a proportion of signaling for each approach angle per individual (n=8). The statistical test was then run on the proportions from each bird. Two birds (one from each study year) never approached their competitor caudally, so these individuals were omitted from the analysis. For this test we report df and p-values.

We tested which pairwise comparisons of approach angle produced significant differences in the Friedman test using a **Conover-Iman test** with a Benjamini-Hochberg correction (Conover and Iman, 1981). This test is appropriate for small sample sizes and few groups. We report pairs of approach angles that accounted for the statistical differences in tail flaring.

Finally, to test the correlation between body mass and the size of a flared tail in morphometric photos, we used a **Spearman Rank correlation** test (Zar, 2005). We report p-values from this test.

### Data availability

Data are currently available in a Dryad repository, and MATLAB and R code for our analyses will be available upon acceptance.

### AI Statement

Machine learning AI was used to digitize anatomical points videos from a training set of points selected by authors (DeepLabCut) (Mathis et al., 2018). These points were quality controlled to ensure coefficients of variance were within acceptable threshold (5%). AI-assisted technologies were used as templates for analysis and visualization code, then further modified and ensured for accuracy of data representation by authors.

### Field observations

As part of a comparative study of hummingbird agonism in the field, we obtained videos for 3D kinematic analysis of free ranging birds. Our field sites were at the Southwestern Research Station (Cochise County, Arizona, USA , “SWRS”) and Santa Rita Lodge (Madera Canyon, Arizona, USA, “Madera”). During this field season (September 1-7, 2024), we recorded video of contests (n = 185) of adult and immature males and females of seven species of hummingbird fighting in the vicinity of artificial feeders: Calliope, Black-Chinned (*Archilocus alexandri*), Rufous (*Selasphorus rufus*), Broad-Billed (*Cynanthus latirostris*), Anna’s (*Calypte anna*), Rivoli’s (*Eugenes fulgens*) and Blue-throated Mountain-gem (*Lampornis clemenciae*). Because these were wild and untagged birds, we were unable to estimate total number of individuals within each species or repeat interactions between birds within each of our two sites. Viewing the videos, we recorded each species involved in the intra or interspecific interaction as well as absence/presence of tail flaring. For this purpose, we used the same criteria used in our lab study of qualitative descriptions of contests (see above) where minimum tail angle which qualifies as flaring to be approximately 30° and the flaring had to be sustained for at least 5 ms. Because active fighting attracted our attention when triggering video recordings, it is likely our samples were biased towards more observable agonism (i.e. accompanied by vocalizations, high-speed or ascending flight), but we attempted to record any agonistic encounter, regardless of the presence of tail flaring.

## Results

### Lab Study

Tail flaring angles among male Calliope hummingbirds during lab trials were different between take-off (−14.1± 7.0 °), and landing (−14.2± 9.1°) versus during competition (when birds won or secured the high perch after a contest: 46.6 ± 41.1 °, and when losing 25.7 ± 42.4 °, example: Video S1). The differences between tail angles during takeoff and competition was significant (Fig. 2, Wilcoxson signed rank test, V = 0, n =10, p<0.001) with a strong effect size (rank-biserial correlation, r=0.89). There was no statistical difference between mean tail angles during takeoff and landing (Wilcoxon signed rank test, V = 26, n =10, p=0.92). There was a statistical effect of winning (secured the high perch) or losing on tail flaring angles (Wilcoxon signed rank test, V = 49, n =10, p=0.01) with a large effect size (rank-biserial correlation, r=0.693).

**Figure 2.**
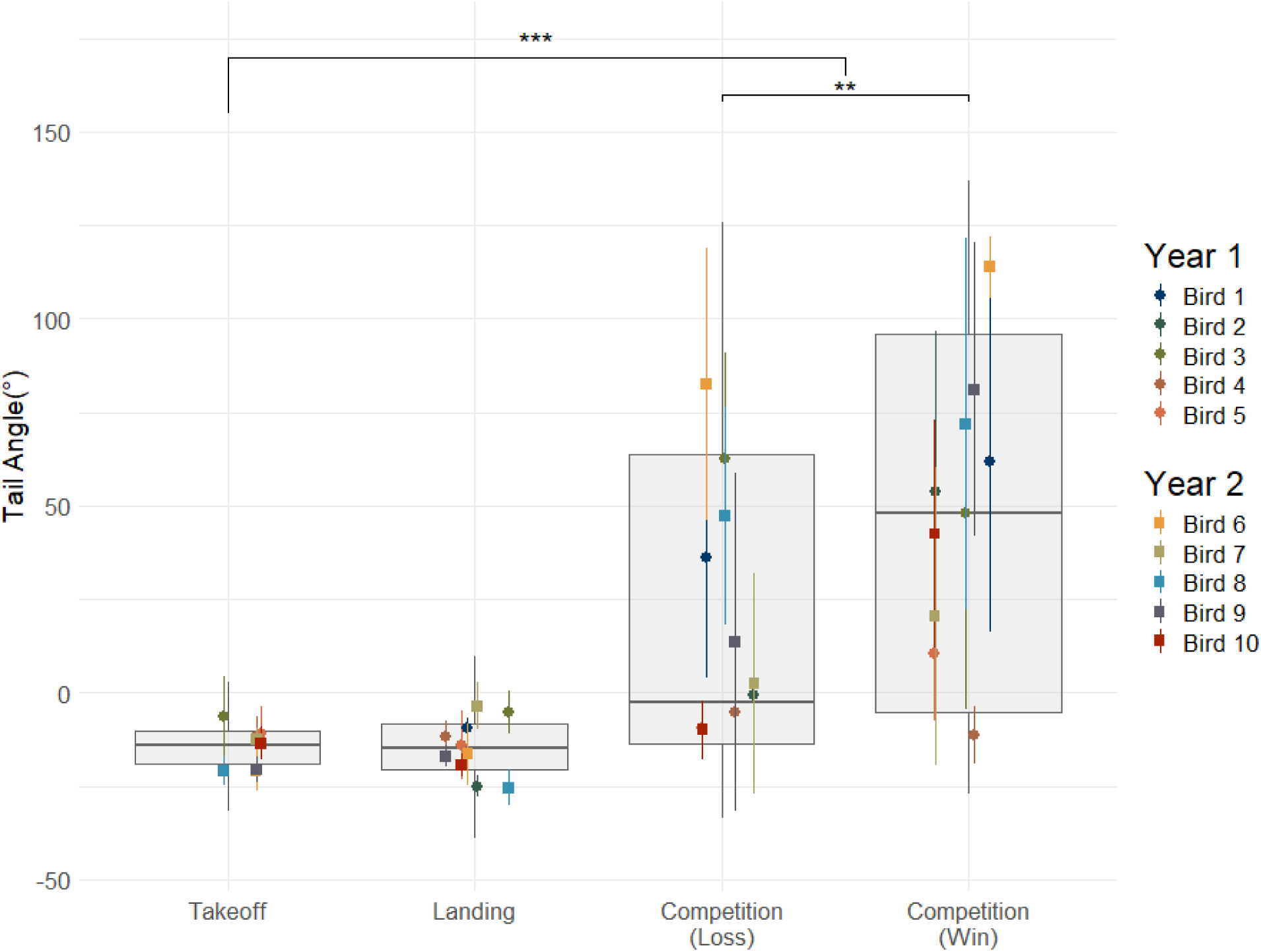
Tail flare angles of Calliope hummingbird, *Selasphorus calliope* (n=10) during agonistic encounters were greatest during competition in a laboratory setting. Individual mean±sd represented as point and whiskers for each condition: solitary takeoffs or landings, compared to contests, which have been split based on contest outcome (win: secured high perch, lose: retreat). Box plots are interquartile range and median values, while vertical lines are minimum and maximum values from all samples. Brackets represent p-values from Wilcoxon Signed-Rank tests performed using n=10, ** : p = 0.01, ***: p <0.001. Birds were numbered based on the order in which they were trapped.

Wilcoxon signed rank tests represent statistical differences between median tail flare angles, yet the variance in tail angle across treatments was compelling to explore. Frequency distributions of tail flaring angles, pooled from all individuals, revealed that there was a greater range of tail flare angles during competition than in solitary conditions (Fig. 3). During takeoff and landing, tail angles distributions were centered at values <0°. Competition had a peak of tail angles below 0°, yet much of the distribution was greater than 50° with some values over 100°. Birds who won contests also had a greater count of tail flare angles at values over 100° when compared with the frequency distribution of losers.

**Figure 3.**
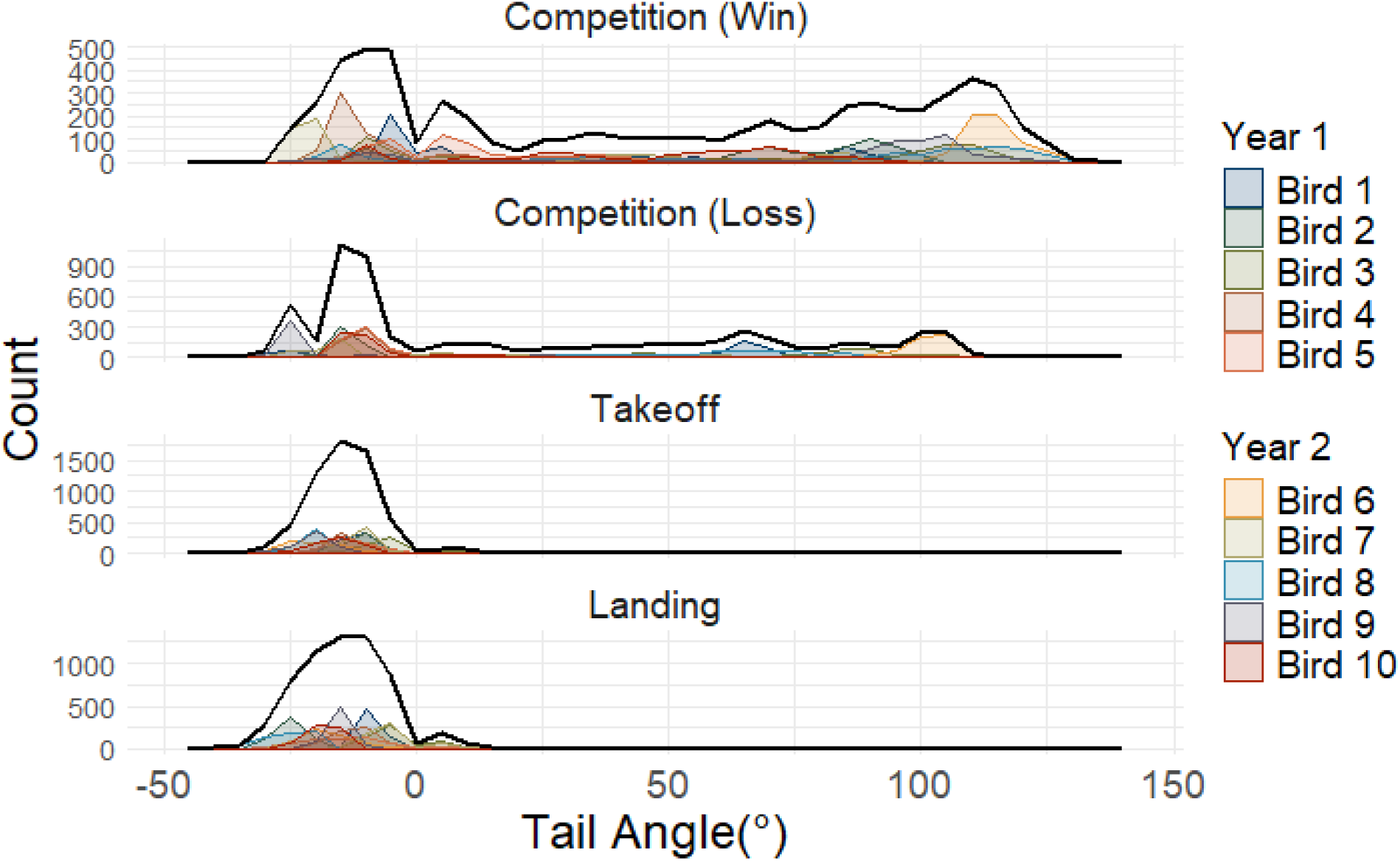
Frequency distributions of tail angles for male Calliope hummingbirds (*Selasphorus calliope, n = 10 birds, pooled data with n = 612 samples from each video per bird*) during landing, takeoff, and winning or losing of a contest. Counts of tail angles are presented for each bird individually, as well as pooled counts for each flight condition. Black lines represent sums of frequencies among birds.

Competing males appeared to exhibit “call and response” of tail flaring. For example, Figure 4 presents tail angles during a fight (595 ms in duration) between a winning bird (green line) who was initially in the air and a bird (yellow line) on the high (desirable) perch who was ultimately displaced. At 63 ms into the encounter, the winning bird flared its tail from 2.6 deg to 120 deg (i in Fig. 3). There was a 50 ms lag before the losing bird started to flare its tail from 1 to 66 deg (ii). The losing bird departed from the perch at 280 ms (iii) and maintained a tail angle averaging 68 deg, much greater than mean tail angles during solitary takeoff (see above). At 400 ms when the losing bird began to retreat in the air (iv), it reduced its tail angle from 71 to 9 deg.

**Figure 4.**
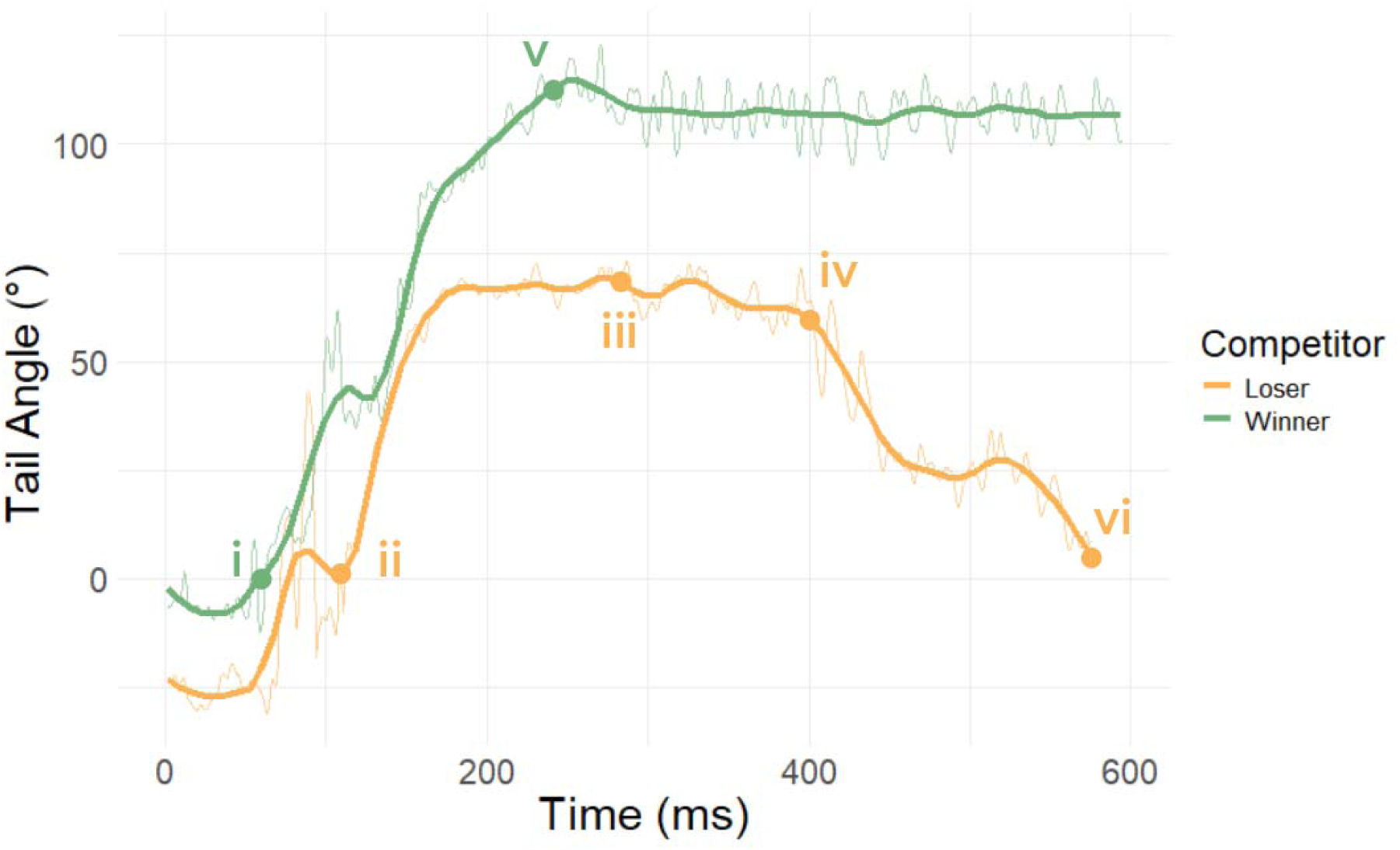
Time series revealing “call-and-response” of angles of tail flaring of two male Calliope hummingbirds (*Selasphorus calliope*) fighting over a desired resource (high perch). The winning bird (green) initiates the displacement by flaring its tail starting at (i), and the occupant of the perch (yellow) responds after lag of 50 ms (ii). The losing bird leaves the perch (iii) and eventually retreats (iv). The winning bird maintains tail flare from (v) to the end of the fight when the loser flies out of the interrogation volume of the video cameras (vi). Fine lines represent raw data smoothed over 5 points (2.5 ms) and bold lines represent these data further smoothed using a LOESS formula.

Throughout the interval from 235 ms (v) to the end of the fight, the winning bird maintained a tail angle of 95 to 124 deg. We considered the fight to end when the losing bird retreated outside the interrogation volume of our cameras (vi).

Our qualitative analysis of tail flaring during approaches toward a bird on the high perch in the flight arena provided further support that tail flaring by the approaching bird was directed toward the perched competitor. There was a significant association between the angle from which a bird was approaching and whether their tail was flared (Friedman test (df=2), p-value=0.002). Post hoc tests revealed significant differences in tail flaring between caudal approaches and front approaches (Conover test with a Benjamini-Hochberg correction, p=<0.001) and between caudal approaches and side approaches (p=<0.001). In other words, every bird flared their tail significantly less often when approaching from behind their opponent.

Tail flaring appeared to increase projected frontal surface area of a competitive male (Fig. 5A,B). Assuming a fixed angle of tail spread of 110° and a vertical upright orientation of the body and tail, our morphometrics revealed that total projected frontal area was significantly greater by 40.3±11.3%, than when tails were not flared (Wilcoxon signed-rank test V=0, p-value < 0.001, r =0.886). This increase in tail area was not accounted for by the birds’ body mass as there was a weak and insignificant correlation between total projected area and body mass (Spearman rank correlation, S =198, rho = −0.2, p =0.584).

**Figure 5.**
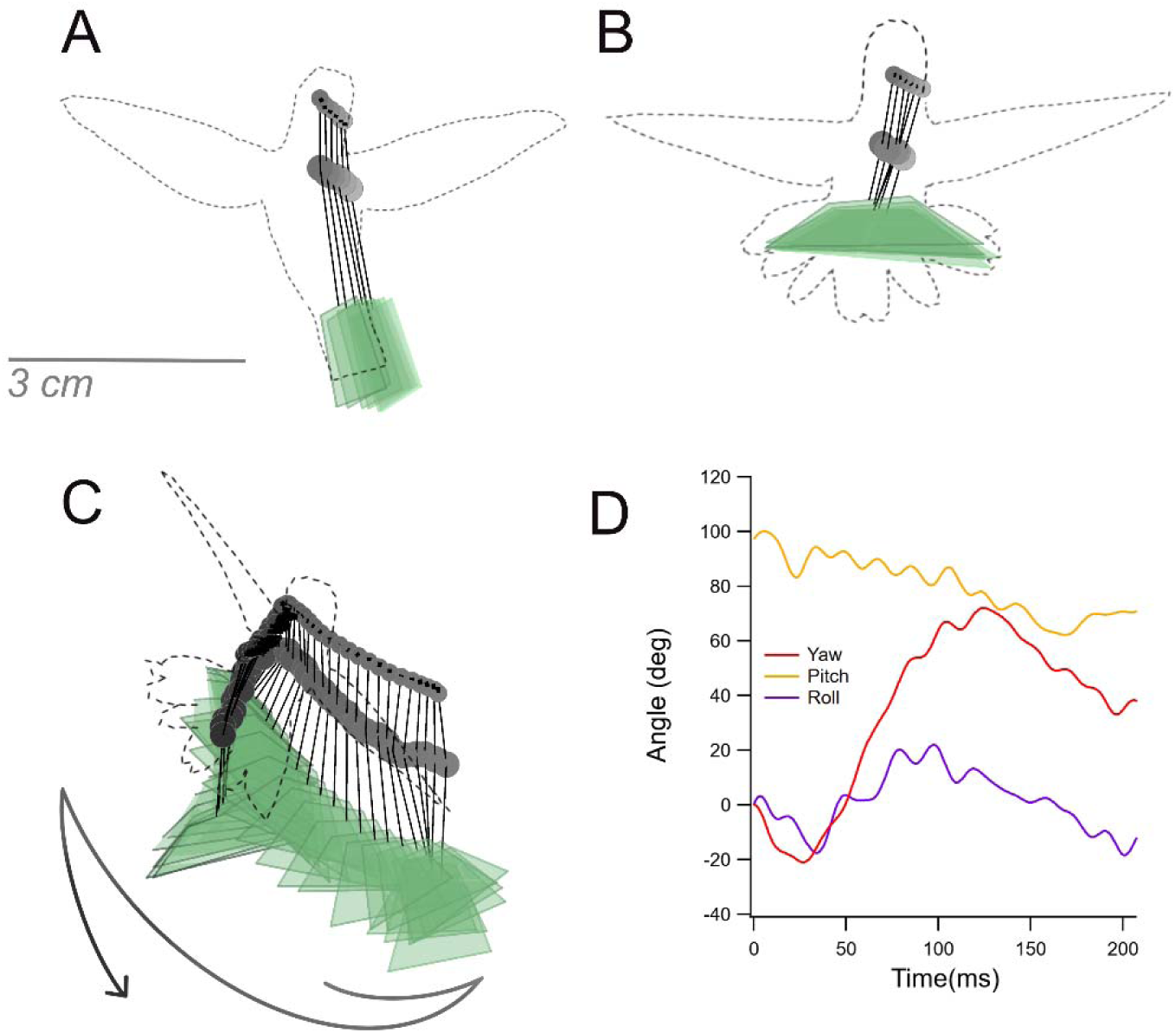
Projected area of the combined body and tail increased when birds flared their tails and was further augmented by “waggle” body movement. Three phases of a display performed by a male Calliope hummingbird (*Selasphorus calliope*) in flight while approaching and intimidating a perched bird. A) tail not flared B) tail flared and C) during a “waggle” where tail flaring was paired with linear translations and body rotations. Positions in A-C sampled every 200 ms. Dashed lines represent body outline at end of display. Solid black lines = beak, green = plane of tail, filled circles = head and center of anterior spine, with shading intensity increasing with time during the display. D) angles of yaw, pitch and roll during the waggle in C (continuously sampled at 2000 Hz).

In addition to the spread of the tail, hummingbirds exhibited body movements with tail flaring in what we here describe as a “waggle” (Fig. 6C) (Hurly et al., 2001). This waggle occurred in ∼44% of encounters that involved tail flaring. One bird who spent the least time on the high perch never performed this behavior and thus this analysis was for n=9 birds. The body movements included linear translations and rotations about all three axes (pitch, roll and yaw, Fig. S1). Among the nine birds who exhibited waggles, average tail angle was 57.9 ± 23° and average pitch angle of the body was 96.4 ±9.8°, meaning the bird was tilted dorsally slightly beyond vertical. Linear translations were 9.5 ± 3.9 cm and 3.7 ± 2.4 cm in the horizontal and vertical directions, respectively. Range of angles about each body axis were 62.2 ± 26.7° for pitch, 55.1 ± 19.2° for yaw and 74.0 ± 31.2° for roll.

**Figure 6.**
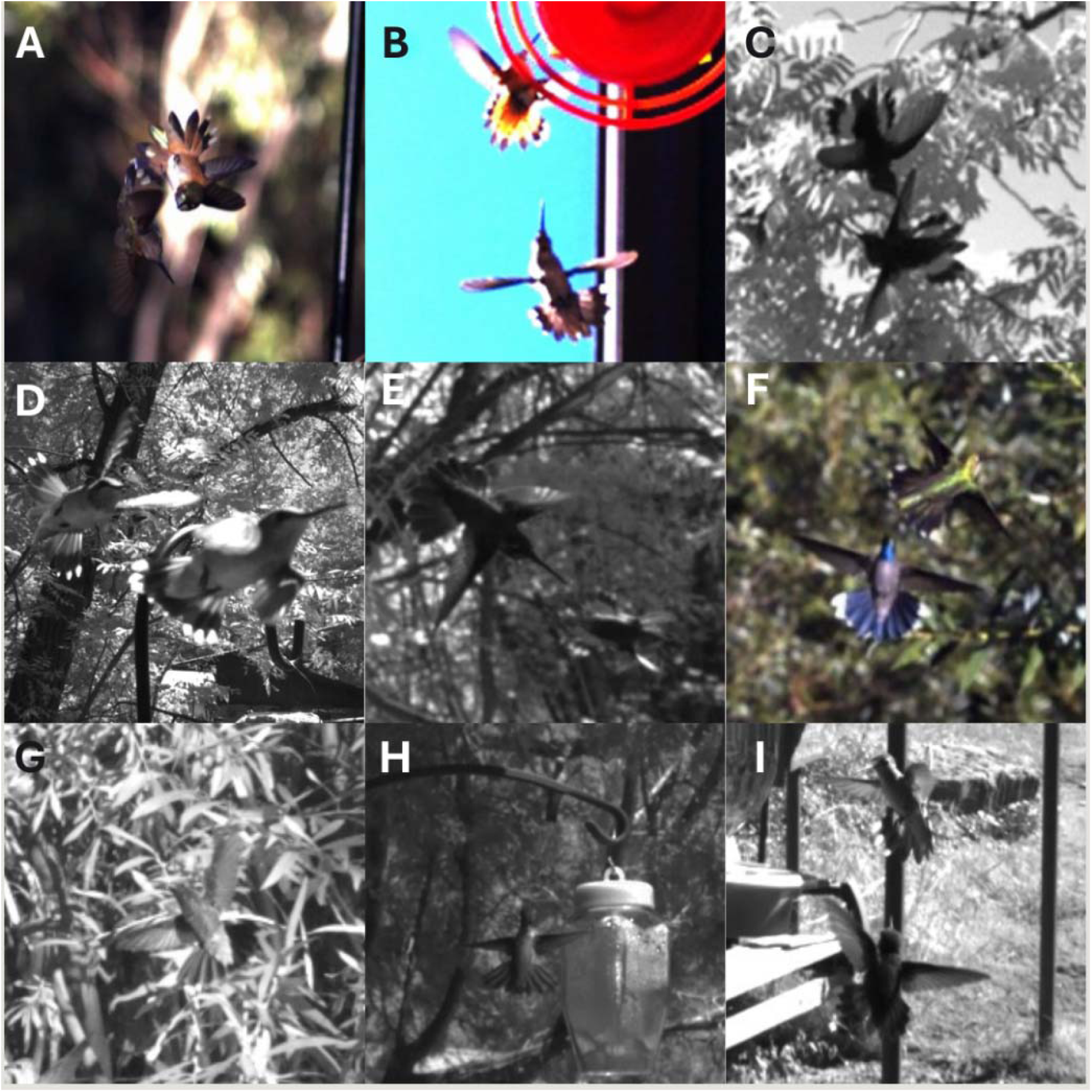
Compilation of flared tails in wild hummingbirds engaged in agonistic encounters in Arizona, USA. A) Rufous (*Selasphorus rufus*, right), chasing Black-chinned (*Archilochus alexandri*, left), B) Rufous (top) and Black-chinned in contest at feeder (red disc), C) two Rufous in ascending fight, D) Black-chinned (left) chasing conspecific, E) Rivoli’s (*Eugenes fulgens*) chasing a Rufous, F) Blue-throated Mountain-gem (*Lampornis clemenciae*) intimidating a Black-chinned, G) Broad-billed (*Cynanthus latirostris*) with flared tail directed at a bird out of frame, H) dorsal view of Anna’s (*Calypte anna)* facing conspecific, I) Calliope (*Selasphorus calliope*) in the foreground flaring tail towards a Black-chinned at a feeder.

We present an example of a bird engaged in tail flaring with waggle display (Fig. 5). Throughout this display, average tail angle was 89.1° and average pitch angle of the body was 80.3°. The bird translated 2.81 cm horizontally and 1.96 cm vertically. He rotated through an approximately symmetrical yaw of 93.1°. Roll angles occupied a 41.3° range and pitch changed angle by 37.8° (Fig. 5D). The waggle maneuver increased the projected area swept by the tail to be 6.43 cm^2^ (Fig. 5C). The equivalent projected areas for his unflared and flared tail were 1.32 cm^2^ and 1.52 cm^2^, respectively (Fig. 5A,B).

### Field study

During field study in Arizona, seven species of hummingbirds (males and females, adults and immatures) in two different locations frequently exhibited agonistic interspecific and intraspecific interactions (Table 1; Fig. 6). We recorded 370 instances of a bird engaged in a fight (note 2 birds per fight), tail flaring was present in 352 (95%) of the instances. The species with the least prevalence of flaring was the broad-billed hummingbird, who flared their tails in 84% of interactions. Our results indicate that tail flaring was routinely used during the behavioral context of inter- and intraspecific agonistic encounters.

**Table 1.** Tail flaring activity in seven species of free-ranging hummingbirds in Arizona (Madera Canyon and Southwestern Research Station). Intra- and interspecific competitions are pooled, and percentages are species specific.

| Species | Sites | Total Contests | Contests with Flaring (%) |
| --- | --- | --- | --- |
| Anna's (*Calypte anna*) | Madera | 8 | 100% |
| Blue-throated mountain-gem (*Lampornis clemenciae*) | Madera, SWRS | 58 | 98.3% |
| Rufous (*Selasphorus rufus*) | Madera, SWRS | 106 | 98.1% |
| Black-chinned (*Archilochus alexandri*) | Madera, SWRS | 159 | 93.7% |
| Calliope (*Selasphorus calliope*) | SWRS | 11 | 90.9% |
| Rivoli's (*Eugenes fulgens*) | Madera, SWRS | 9 | 88.9% |
| Broad-billed (*Cynanthus latirostris*) | Madera, SWRS | 19 | 84.2% |

## Discussion

We found greater tail flaring angles in male Calliope hummingbirds during competition trials than in other contexts. These results support our hypothesis that the posture of hummingbirds’ tails varies between locomotor actions (takeoff and landing) and agonistic competition, suggesting that this flaring is used most often in a social contest. Greater tail flare angles were correlated with winners, while lower tail flare angles were associated with males who lose (Figs. 2 and 3). Hummingbirds did not flare their tails as frequently when out of the view of their competitor. While our sample sizes for detailed analyses in the lab were modest and only from Calliope hummingbirds, the conclusion that tail flaring is associated with competition was corroborated by high prevalence (>90%) of tail flaring during interactions among seven species of wild hummingbird including juveniles and females (Fig. 6, Table 1). The maximum angles of tail flaring during competition appeared similar to those during rapid escape performance in Calliope hummingbirds (Anwar et al., 2024) and four other species of hummingbirds (Haque et al., 2023b). However, the duration of tail flaring during agonism lasted much longer than during escape (often over 1 second compared to 120ms, for example 600 ms in Fig. 5).

Our results suggest that flaring of the tail is used for communication with other hummingbirds and this form of communication is consistent with the hummingbird visual system. Tail flaring and waggling (Fig. 5) can be easily perceived by other hummingbirds based on current understanding of the hummingbird visual system. Hummingbirds are near-sighted (equivalent of ∼20/100) (Altshuler and Wylie, 2020) with a region of binocular vision (∼29°) by the bill tip, monocular vision on the sides of the head, and a blind spot at the back of the head (∼22°) (Tyrrell et al., 2018). This blind spot probably accounts for the observation that birds infrequently flared their tail in caudal approaches, given this is outside of the visual field and communication with a visual signal is not effective here.

We propose that the use of tail flaring and waggle displays are attempts to intimidate opponents via a looming stimulus. Flaring by itself increased the projected body size presented to a competitor, but the addition of waggling further increased this size (Fig. 5). These two behaviors together present a looming stimulus to a competitor, and looming in many taxa elicits an evasive response (Peek and Card, 2016). Waggling likely causes an individual to spend time in both the monocular and binocular visual field of their opponent. Looming stimuli depend on the fractional rate of change of an approaching item on the retina, and the efficacy of looming depends on estimates of time-to-contact (TTC) (Peek and Card, 2016). TTC is best discriminated when an opponent is in the binocular visual field (Gray and Regan, 2004), but the increased ganglion cell density and invagination of the central retina (fovea centralis) in hummingbirds (Tyrrell et al., 2018) allow for spatial acuity, and probably accurate TTC estimates, even in the monocular region. That written, in humans, the addition of horizontal motion to a stimulus reduces TTC accuracy (Calabro et al., 2011). Tail flaring with added waggling during approaches from the front or side (Fig. 5) may introduce uncertainty in TTC estimates and may augment the intimidation effect.

Flaring of the tail during competition and its potential role for intimidation align with current definitions of signals used for animal communication. An incoming bird who initiated a contest (Figs. 4 and 5) flared its tail, which may communicate an intention of attack with force if its opponent does not retreat (Bradbury and Vehrencamp, 2011). The continued use of tail flaring throughout the interaction could also mean this is a signal that directly communicates some aspect of the quality of the opponent. If body size is an important quality to signal, as it is in other taxa (Brown et al., 2005; Guilford and Dawkins, 1995; Umbers et al., 2012; Vieira and Peixoto, 2013), we might expect the use of a signal that is constrained by a physical property of the signaler where only the largest individuals have the largest signals (i.e., an index signal) (Bradbury and Vehrencamp, 2011). If true, then we would expect large individuals to flare their tails the most widely. This was not supported by our data, as body mass was not correlated with projected frontal area when the body had a flared tail. Instead, our results suggest that there is a unique distribution of maximal tail flare angles in individuals that win the high perch (Fig. 3), but maximal angles were not correlated with body mass. Of the highest quantile (80^TH^) of tail flare angles that we measured, over 15% of maximal angles were exhibited by the bird with the *smallest* body mass (Bird 3 in Figs. 2 and 3 and “winner” in Fig. 4). This individual also spent the most time on the desirable high perch (Gaffney, 2017) via successful displacements.

If the hummingbird tail is a signal of individual fighting ability, or resource holding potential (RHP) (Parker, 1974), then flaring for communication would help explain the rarity of lethality and injury in field-observations of hummingbird fighting (Baltosser and Russell, 2020; Evens and Harper, 2020). Fights are not only non-lethal, but often non-physical. Bird species that have evolved for increased aerial lifestyles often have reduced weapons (e.g. bony spurs) (Menezes and Palaoro, 2022). Many bird taxa have evolved alternative, often multi-modal, auditory and visual signals for use in important, non-physical stages of contests (Wiens and Tuschhoff, 2020) . As hummingbirds are aerial specialists who could spend over half of their day on the wing (Elting et al., 2026), it is likely they use signaling to resolve most contests before escalation to physical contact. Effective signaling requires the signal quality to reflect individual quality (RHP). When opponents can estimate the RHP of a rival from the quality of a signal, it often pays the weaker competitor to cede the contest before it escalates to dangerous battle (Smith and Harper, 2003; Zahavi, 1975). Agonistic signals have been well studied in a diversity of animal contests (Brown et al., 2005; Emlen et al., 2012; Green and Patek, 2018; Vieira and Peixoto, 2013; Yasuda et al., 2011), but little is known about this sort of signaling in hummingbirds. Tail flaring has been described in fights of other hummingbirds (Hurly et al., 2001; Williamson, 2020) and we show here that it is used extensively during fighting in seven species in the field, and in male Calliope hummingbirds in our competition arena.

### Future directions to test if tail flaring is an honest signal

If flaring does signal individual quality, we predict that this is due to the metabolic cost of flaring the tail, not the cost of growing or carrying it. The energetic cost for developing tail feathers in minimal (Buttemer et al., 2020) and the tail only contributes marginally to body mass and aerodynamic drag (Clark and Dudley, 2009). These costs are energetically inexpensive when compared with signals of male quality in other taxa (e.g. horns, antlers, overall body size (Emlen, 2008)). Tail feathers are molted seasonally and are easily shed in response to predatory attack (Farner and King, 2013). The ability of tail flaring to communicate individual quality may instead rely on the energy used by muscles to flare the tail. Overall muscle quality is important in combat, allowing for an individual to chase faster (Sholtis et al., 2015) or be more agile in fighting. Muscular contractions by the *bulbi retricium* muscle (Gatesy and Dial, 1996) are responsible for the flaring of the tail. Tail flaring during contests can last up to several seconds, and the energetic cost of muscle contraction should be proportional to the duration of contraction and the number of motor units recruited (Taylor et al., 1980). If individuals with the highest RHP also flare their tails at the greatest angles or for the longest periods of time, this could be signaling an individual’s overall muscle quality. If flaring adversely affects the aerodynamics of hovering through wake interactions, (Su et al., 2012), alteration of body pitch, or changes in wing trajectories, perhaps this can only be accomplished by individuals with the highest RHP. Tail flaring that is accompanied by waggles may further accentuate RHP if waggling increases metabolic costs beyond steady hovering (Groom et al., 2018).

To understand signaling of the tail, whether it communicates quality and functions to modulate contest escalation within hummingbird agonism, it will be necessary to study tail flaring in relation to other phenotypic traits including morphology, behavior, and physiology. Traits that are already understood to improve flight performance in hummingbirds include marginal power availability which correlates with agility (Altshuler et al., 2004; Dakin et al., 2018; Segre et al., 2015). In other bird species, individual aggression and overall body condition (Careau et al., 2008; Dolnik and Hoi, 2010), including bilateral symmetry (Møller, 1992) have explained differences in male competitive and reproductive success. Variable time budgets may be associated with aggression (Ewald and Bransfield, 1987), and if muscular activity from flaring of the tail comes at a metabolic cost it could be important to consider daily energy expenditure on these tasks (Shankar et al., 2019). To rigorously test whether tail flaring correlates with individual quality, it will be necessary to carefully investigate phenotypes at the individual level in behavioral experiments that control for residency effects (Alcock and Bailey, 1997; Mendiola-Islas et al., 2016) and repeat interactions. Together, these would further our understanding of the evolution of dominance in contests, phenotypes associated with competition success and tail flaring as a signal in hummingbird agonism.

### Alternative hypotheses

Flaring of the tail seemed important for communication during agonism, but this flaring may also serve an aerodynamic function during contests. Tails that were flared during competition exhibited small oscillations (∼15°) in angle approximately in phase with the wingbeat cycle (faint lines in Fig. 4). In-phase flaring in the passerine species, the warbling white-eye (*Zosterops japonicus*), is used to mitigate excess body pitch (Su et al., 2012). The wingbeat kinematics of this passerine varies significantly from those of hummingbirds; an asymmetric flexed-wing upstroke of the passerine provides minimal aerodynamic function (Crandell and Tobalske, 2015), while the active upstroke of the hummingbirds’ fully-extended wing generates lift (Warrick et al., 2005). In hummingbirds, the wings dominate the aerodynamics (Haque et al., 2023a), but high-frequency of modulation of the tail in our work suggests the tail could be providing minor pitch stabilization via aerodynamic forces. In addition, it is possible that the inertia of the tails of hummingbirds could function as vibration absorbers and dampers to suppress the body oscillations induced by wing flapping (Haque et al., 2023b).

Even during the dramatic maneuvers of escape flight from a looming threat, the tail contribution to overall torque is minimal compared to the contribution of the wings (Nasirul Haque et al., 2023). One species of hummingbird (black-chinned hummingbird, *Archilocus alexandri*) is known for their exaggerated and diagnostic (Sibley, 2014) “pumping” of the tail during slow flight. Females and males of this species modulate tail flare angles and angle of the tail plane relative to the body plane. This species additionally flared their tail in ∼94% of interactions in our field study (Table 1). Further study of this species would be a promising direction to test for an evolutionary tradeoff between flaring of the tail for slow flight aerodynamic benefits versus communication in hummingbirds.

It is also possible that tail flaring is a byproduct of the intensity and arousal that occurs during competitive interactions or is part of a generalized aggressive phenotype. When assigned a difficult task, human infants not only perform the task with the assigned hand, but movement also occurs with the free hand (Soska et al., 2012). This motor overflow is characterized by an intended movement during strenuous activity being accompanied by a secondary movement.

Hovering is the most costly form of flight (Tobalske et al., 2003), so during slow flight the strenuous demand of activating wing muscles could be accompanied by secondary tail muscle activation. A second alternative hypothesis would be that the flaring of the tail is part of a generalized aggressive signal. During courtship, male Broad-tailed hummingbirds (*Selasphorus platycercus*) synchronize generation of sounds from the tail with display of their iridescent gorget to females (Hogan and Stoddard, 2018). Given that many birds possess multiple signaling modalities (Menezes and Palaoro, 2022), synchronization of signals may be common among birds. Thus, flaring of hummingbird tails may serve a communication role among a suite of traits that form an aggressive signal instead of an independent signal that varies with individual quality.

## Conclusions

Tail flaring was regularly used by male Calliope hummingbirds during competition trials in an indoor competition arena as well as among free-ranging hummingbirds in seven species at two different field sites. The angles of tail flaring were different between solitary maneuvers and during competition, with statistical differences between birds that won a desired resource (high perch) and those that did not. The use of tail flaring was variable and did not associate with body size, therefore we propose that it and concurrent waggle maneuvers may serve as signals of aggressive intent in male-male fighting in hummingbirds. Our field observations indicate such signaling is also performed by females. Testing for correlations between tail flaring and an individual’s quality (Resource Holding Potential, RHP) may be revealed through further study of the metabolic energy requirements and aerodynamics of aerial competition that include tail flaring.

## Acknowledgements

We thank K. Dial, B. and D. Krieske, and the friendly staff at Southwestern Research Station and Santa Rita Lodge, Madera Canyon, for access to their land for hummingbird trapping and videography.

## Author Contributions

RLE: project conceptualization, data curation, investigation, methodology, formal analysis, visualization, writing-original draft. MZA: data curation, investigation, formal analysis, writing-review and editing. DRP: data curation, investigation, methodology, writing-review and editing. BC: project conceptualization, funding acquisition, writing-review and editing. HL: project conceptualization, funding acquisition, writing-review and editing. BWT: project conceptualization, funding acquisition, data curation, investigation, methodology, formal analysis, supervision, writing-review and editing.

## Data and resource availability

Raw data associated with this manuscript are published on a dryad digital repository: 10.5061/dryad.dz08kpsbj. R scripts for data analysis will be available on GitHub upon acceptance.

