## Supplemental File for "Tail Flaring During Agonism in Hummingbirds"

### Supplemental Information

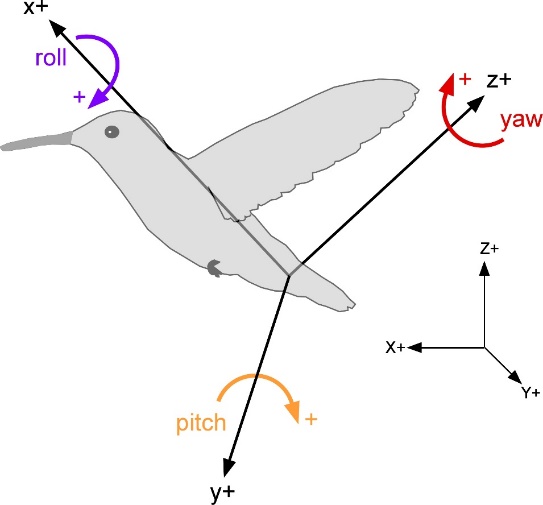

**Figure S1.** Global (X, Y, Z) and bird-centered (x, y, z) coordinate systems.

[*Link*](https://drive.google.com/file/d/1DNgBTiqivEYRNqAiEeN25VXDV20aMFSr/view?usp=sharing)

**Video S1**: Multiple views of tail flaring by a male calliope hummingbird (*Selasphorus calliope*) displaying to a male on the high perch (desired resource) in our contest arena. The displaying male (Bird 3 in the main manuscript) regularly displaced competitors and secured the high perch in competitions. Original video sampled at 2000 frames s^-1^, with playback at 200 frames s^-1^.

**Table S1.** Coefficients of variance (CV) in length of the bill and width of the rump digitized from male calliope hummingbirds (*Selasphorus calliope*). We include each combination of behavior and individual in our experiments in the competition arena. Rump widths changed as the hummingbirds flared their tails, therefore, we report the Pearson correlation coefficient between rump width and tail angle. In our methods of quality control, we ensured CV < 5% for fixed length of a given bird’s bill.

| **Bird number** | **Treatment & Outcome** | **Bill CV** | **Rump CV** | **Pearson Correlation Coefficient** |
| --- | --- | --- | --- | --- |
| 1 | Competition (Win) | 2.06 | 27.34 | 0.90 |
| 1 | Competition (Loss) | 3.61 | 14.04 | 0.46 |
| 1 | Landing | 4.35 | 6.39 | -0.43 |
| 1 | Takeoff | 3.55 | 6.00 | -0.41 |
| 2 | Competition (Loss) | 2.25 | 13.13 | 0.48 |
| 2 | Competition (Win) | 2.04 | 15.73 | 0.96 |
| 2 | Landing | 2.51 | 6.03 | -0.47 |
| 2 | Takeoff | 4.07 | 7.50 | 0.76 |
| 3 | Competition (Loss) | 2.65 | 10.80 | 0.79 |
| 3 | Competition (Win) | 2.40 | 13.12 | 0.85 |
| 3 | Takeoff | 4.01 | 8.08 | -0.31 |
| 3 | Landing | 3.79 | 4.92 | -0.52 |
| 4 | Competition (Loss) | 3.35 | 14.62 | 0.83 |
| 4 | Competition (Win) | 2.18 | 10.95 | 0.30 |
| 4 | Landing | 3.89 | 10.31 | -0.83 |
| 4 | Takeoff | 3.98 | 18.58 | -0.55 |
| 5 | Competition (Loss) | 2.37 | 8.07 | 0.02 |
| 5 | Competition (Win) | 1.78 | 14.66 | 0.69 |
| 5 | Takeoff | 3.91 | 11.46 | -0.51 |
| 5 | Landing | 4.43 | 15.42 | -0.74 |
| 6 | Competition (Loss) | 1.6613 | 10.9434 | 0.8049 |
| 6 | Competition (Win) | 4.3344 | 7.8316 | -0.046 |
| 6 | Takeoff | 4.5824 | 9.4907 | -0.7676 |
| 6 | Landing | 3.0602 | 6.0064 | -0.0267 |
| 7 | Competition (Loss) | 3.3796 | 15.17 | 0.7206 |
| 7 | Competition (Win) | 2.1124 | 9.0426 | 0.8183 |
| 7 | Takeoff | 1.8674 | 7.5372 | -0.0815 |
| 7 | Landing | 2.6583 | 2.7069 | -0.0232 |
| 8 | Competition (Loss) | 4.7884 | 11.4611 | 0.7367 |
| 8 | Competition (Win) | 2.3298 | 17.137 | 0.8274 |
| 8 | Takeoff | 2.0092 | 5.2354 | -0.3595 |
| 8 | Landing | 2.4427 | 6.6023 | -0.6848 |
| 9 | Competition (Loss) | 2.5579 | 9.9052 | 0.801 |
| 9 | Competition (Win) | 2.9845 | 18.1459 | 0.8036 |
| 9 | Takeoff | 1.9726 | 8.5034 | -0.0759 |
| 9 | Landing | 1.8754 | 7.0543 | -0.8112 |
| 10 | Competition (Loss) | 2.5836 | 5.5003 | 0.3586 |
| 10 | Competition (Win) | 3.9619 | 12.9912 | 0.7308 |
| 10 | Takeoff | 3.5523 | 7.2384 | -0.6896 |
| 10 | Landing | 1.8598 | 6.2767 | -0.5934 |
